# Chronic High Intensity Interval Training (HIIT) exercise in adolescent rat’s result in cocaine place aversion and ΔFosB induction

**DOI:** 10.1101/2024.12.10.627707

**Authors:** Nikki Hammond, Nabeel Rahman, Sam Zhan, Mark S Gold, Kenneth Blum, Teresa Quattrin, Yun Young Yim, Eric J Nestler, Panayotis K. Thanos

**Affiliations:** Behavioral Neuropharmacology and Neuroimaging Laboratory on Addictions (BNNLA), Research Institute on Addictions, Department of Pharmacology and Toxicology, Jacobs School of Medicine and Biomedical Sciences, University at Buffalo, Buffalo, New York, United States of America; New York College of Podiatric Medicine, New York, New York, United States of America; Department of Psychology, University at Buffalo, Buffalo, New York, United States of America; Nash Family Department of Neuroscience and Friedman Brain Institute, Icahn School of Medicine at Mount Sinai, New York, New York, United States of America; Department of Psychiatry, Washington University School of Medicine, St. Louis, Missouri, United States of America; Center for Sports, Exercise, and Mental Health, Western University of Health Sciences, Pomona, California, United States of America; Department of Molecular Biology, Adelson School of Medicine, Ariel University, Ariel, Israel; UBMD Pediatrics, JR Oishei Children’s Hospital, University at Buffalo, Buffalo, New York, United States of America

**Keywords:** Exercise, Cocaine Conditioned Place Preference, HIIT, Cocaine Substance abuse, ΔFosB, Dopamine

## Abstract

High-Intensity Interval Training (HIIT) is a form of exercise that has been greatly popularized over the past few years for its many health benefits. Similar to other forms of exercise, HIIT may be beneficial in the prevention of substance use behaviors; however, the extent to which HIIT can impact the reinforcing effects of drugs of abuse during adolescence has not been fully evaluated. Here, we assess the effects of HIIT during adolescence on subsequent cocaine conditioned place preference (CPP) in male Lewis rats. The HIIT exercise exposed rats ran on a treadmill for 30 minutes daily (ten three-minute cycles) for six weeks with progressive speed-increased up to 0.8 mph (21.5m/min), while the sedentary rats remained in their home cage. Following the six weeks of exercise, rats were tested for cocaine (25 mg/kg) CPP. Following completion of the behavior test ΔFosB levels were measured in the brain. Results showed that the HIIT rats showed significantly attenuated place preference (-19%) in their time spent in the cocaine-paired chamber compared to the sedentary environment rats. In addition, HIIT rats had significantly higher (65%) striatum ΔFosB levels compared to the sedentary rats. Our findings show that HIIT exercise during adolescence could be protective against cocaine abuse which may be mediated by an increase in ΔFosB. This finding has important clinical implications with respect to exercise mediated protection against substance misuse and abuse. Future studies will examine this effect in females as well as the potential underlying mechanisms.

## 1. Introduction

In 2019, 23 million individuals ages 12 and up were reported to suffer from a substance abuse disorder [1]. In particular, one of the most frequently abused drugs today is cocaine, ranking second in illegal drug use after cannabis globally [2]. Cocaine primarily increases synaptic dopamine levels by inhibiting dopamine reuptake. Chronic cocaine use results in increased neuronal dendritic branching and spine density in the nucleus accumbens and prefrontal cortex, which is thought to increase the incentive behind drug use [3]. Cocaine abuse also alters the mesolimbic reward pathway in the brain, in part through increased ΔFosB expression in the nucleus accumbens [4].

Previous research has supported the use of exercise for both the prevention and treatment of substance abuse. Physical activity has been shown to decrease cocaine conditioned place preference (CPP), attenuate cocaine cue-induced reinstatement, and inhibit stress-induced reinstatement of cocaine CPP in rodents [5–7], while also reducing cocaine craving and usage in humans [8].

Different exercise regimens display differing levels of efficacy, including varying therapeutic potential regarding neuropsychiatric diseases [9]. High-intensity interval training (HIIT) has been shown to result in greater improvements in VO_2_ max values compared to moderate-intensity continuous exercise (MICT) [10], lowered insulin resistance and decreased fasting blood glucose levels [11], enhanced cognitive performance and working memory capacity [12]. Inactive people are also more likely to continue exercising under a HIIT regimen than MICT [13]. Previous research has shown that MICT during adolescence attenuated future cocaine place preference behavior in females while blocking in males [7]. The present study examined the impact of HIIT treadmill exercise during adolescence on cocaine preference in male rats.

ΔFosB, a member of Fos family of proteins, has been found to play a significant role in addictive behaviors that are associated with addiction [14–17]. The ability of ΔFosB to increase sensitivity to drugs of abuse and increase drug seeking behavior has led to it being labeled a sustained molecular switch [17–19]. Following chronic exposure to cocaine, ΔFosB—but not any other Fos family protein—accumulates and its expression remains stable for weeks [19–22]. This accumulation of ΔFosB is also seen following chronic cocaine self-administration as well as yoked exposure [21, 22]. In this paper, we measured the level of ΔFosB expression in the striatum following HIIT and subsequent cocaine place preference with the hypothesis that rats exposed to HIIT exercise would have lower levels compared to sedentary controls.

## 2. Materials and Methods

### 2.1 Animals

Male (n = 32) Lewis rats were obtained at 6 weeks of age (Charles River Laboratories Incorporated). Subjects were housed under standard laboratory conditions (22°C ± 2°C; 12-hour reverse light/dark cycle [lights off: 08:00 – 20:00]. Food and water were available ad libitum for the duration of the study. Body weights of all subjects were measured daily. This experiment was conducted in accordance with the National Academy of Sciences Guide for the Care and Use of Laboratory Animals (1996) and University at Buffalo Institutional Animal Care and Use Committee (Protocol Number: 202100079).

### 2.2 Drugs

Cocaine (Sigma-Aldrich, St. Louis, MO, USA) was dissolved in saline at a concentration of 12.5 mg/ml and administered at a dose of 25 mg/kg [7,23].

### 2.3 Apparatus

#### 2.3.1 Treadmill

A custom-made motorized treadmill was used to conduct forced exercise on the experimental rats (Figure 1A). The treadmill was comprised of four Plexiglas running lanes, each with dimensions of 25 in. x 4.5 in. x 21 in. (L x W x H).

**Fig 1.**
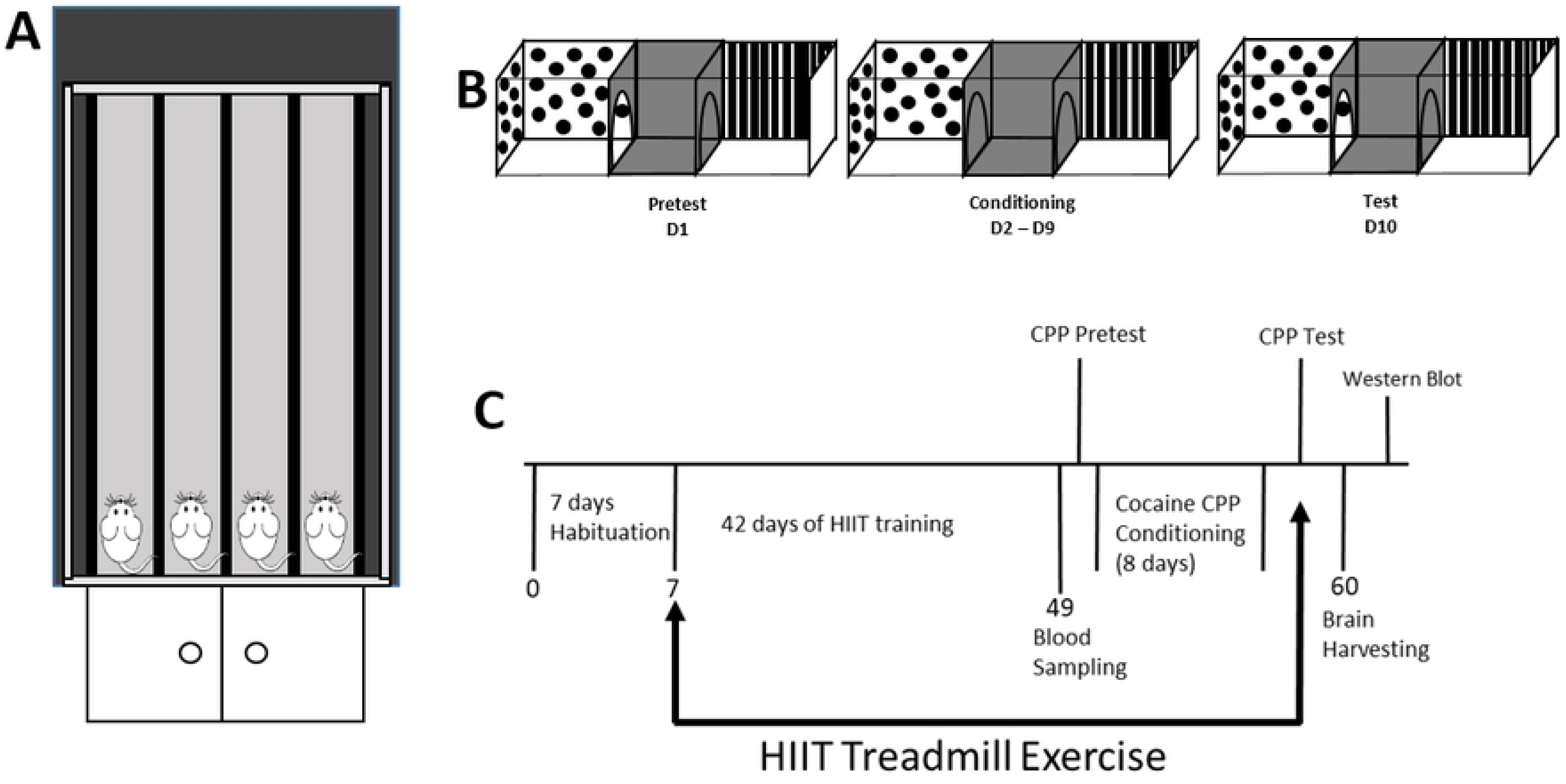
Experimental design. A) The treadmill apparatus used for the HIIT daily exercise included four equally spaced Plexiglas lanes. B) Cocaine conditioned place preference timeline. Pretest (Day 1): Free access given to both chambers; drug paired chamber and preferred chamber determined for each subject. Conditioning (Day 2 – Day 9): Subjects confined to saline or cocaine paired chamber on cocaine conditioning days. Test (Day 10): Free access given to both chambers; time spent in cocaine paired chamber measurement. C) The experiment timeline.

#### 2.3.2 Conditioned Place Preference (CPP) Boxes

The CPP apparatus consisted of two compartments connected by a central corridor (Figure 1B). The compartments each possessed dimensions of 12 in. x 7.5 in. x 8 in. (L x W x H). The central corridor possessed dimensions of 4.75 in. x 8.25 in. x 8.25 in. (L x W x H). One compartment consisted of a perforated stainless-steel floor and black and white striped walls. The other compartment consisted of a smooth floor with black and white polka dot walls. The central corridor consisted of a smooth floor with dark gray walls.

### 2.4 Procedure

#### 2.4.1 Lab Habituation

After arrival to the laboratory rats underwent one week of habituation during which they had daily handling and body weight measurements (Figure 1C).

#### 2.4.2 High Intensity Interval Training (HIIT) Exercise Regimen

Subjects were divided into two groups: a sedentary group (n=16) and a HIIT group (n=16). Sedentary subjects remained in their home cages for the duration of the study. HIIT subjects were exposed to the chronic exercise regimen (see Figure 1C). Exercise was carried out seven days a week for six weeks. Subjects first underwent five days of habituation to the treadmill at a speed of 0.64 km/h (10 m/min) and for a duration of 10 minutes. Subjects then began the HIIT regimen. Each HIIT session lasted for 30 minutes and consisted of ten three-minute exercise cycles. Each exercise cycle consisted of two minutes of active running followed by one minute of sedentary rest. Running speed began at 0.64 km/h (10 m/min) and was increased by 0.16 km/h (2.68 m/min) every five days until the top speed of 1.29 km/h (21.46 m/min) was reached. Exercise then continued at the top speed for the remainder of the exercise regimen. If needed, an air puff at the end of the treadmill lane was used to maintain running. All exercise sessions were performed during the subjects’ dark cycle (08:00 – 12:00).

#### 2.4.3 Cocaine Conditioned Place Preference (Cocaine CPP)

After the chronic exercise regimen had been completed, Cocaine CPP was carried out. The CPP procedure consisted of three phases, Pretest, Conditioning, and Test, and spanned for ten days in total. During Pretest (Day 1) subjects were given free access to the entire CPP apparatus for 15 minutes (See figure 1B). Time spent in each compartment was recorded. The compartment in which more time was spent was defined as the *preferred chamber*, while the compartment in which less time was spent was defined as the *drug-paired chamber*. During Conditioning (Day 2 – Day 9), subjects were given cocaine and saline on an alternating day scheduling, such that cocaine administration was followed by saline administration the following day. Cocaine and saline administration occurred via I.P. injection. Following saline administration, subjects were placed in the *preferred chamber* for 15 minutes. Following cocaine administration, subjects were placed in the *drug-paired chamber* for 15 minutes. Conditioning was carried out for eight days. During Test (Day 10) subjects were once again given free access to the entire CPP apparatus for 15 minutes, as was done during Pretest. Time spent in each compartment was recorded, and Cocaine CPP was determined by comparing time spent in the *drug-paired chamber* on Test Day to time spent in the *drug-paired chamber* during Pretest.

#### 2.4.4 Stress Reactivity Testing (Serum Corticosterone ELISA)

Following the end of the HIIT regimen and prior to CPP, subjects’ blood was obtained to assess serum corticosterone levels via an enzyme-linked immunosorbent assay (ELISA). Blood was obtained via tail vein sampling, which was performed as subjects were under light anesthesia (≈ 2.5% isoflurane). Blood was allowed to clot for 30 minutes and was then centrifuged for 15 minutes at 4°C 3000 RPM. Serum was extracted and then stored at ≈ - 80°C until being assayed in triplicates for corticosterone using an ELISA (IBL TECAN, Charlotte, North Carolina) according to the manufacturer’s instructions. The CORT ELISA was then ran through a BioTek CYTATION1 Imaging reader. The Imaging reader used the BioTek Gen5 Data Analysis Software to determine the absorbance of each well at 450nm.

#### 2.4.5 Tissue Preparation (Western blot sampling)

Rats were euthanized 24 hours after CPP test day (Figure 1C). Briefly, rats were euthanized under deep isoflurane anesthesia (∼3.0%). Brains were harvested quickly, flash frozen in 2-methlybutane, and stored at -80 ֯C. Bilateral 1mm tissue punches were taken from the dorsal and ventral striatum in a cryostat at -20 ֯C. Brain punches were weighed and mashed in a 1:40 initial dilution of Pierce^TM^ IP Lysis Buffer (25 mM Tris-HCl pH 7.4, 150 mM NaCl, 1 mM EDTA, 1% NP-40 and 5% glycerol) with Halt^TM^ Protease Inhibitor Cocktail (100X). Homogenates were then spun in a centrifuge for 30 minutes at 4 ֯C 10,000 RPM. Supernatant was then collected and stored in a -80 ֯C freezer.

#### 2.4.6 Immunoblot Analysis (Western blot)

Aliquots (20 μg of each sample) were ran on an 8-16% polyacrylamide gel for SDS-PAGE and then electrotransferred to nitrocellulose membrane. The blots were blocked twice (30 min per wash) with 0.5% dry milk in TBS-Tween (1X TBS containing 0.1% tween 20) at the room temperature. Then, the blots were washed five times for 5 min each with TBS-Tween (TBST) at room temperature and incubated overnight with FosB anti-rabbit antibody (Cell Signaling, 1:1000) in blocking buffer at 4°C. Next day, the blots were washed five times for 5 min each with TBST and incubated for 1 hr with goat anti-rabbit antibody conjugated to horseradish peroxidase (KPL, 1:10,000) in blocking buffer at room temperature. Then, the blots were washed 3 times for 10min each with TBST, followed by two additional washes with TBS (15 min per wash). Using Western Lightning™ Chemiluminescence Reagent Plus (Perkin-Elmer) and iBright imager (TheromoFisher), Western blots were imaged. For the β-actin (loading control) detection, the blots were stripped using Restore™ plus Western blot stripping buffer and reblotted as stated above. β-actin anti-mouse antibody (Cell Signaling, 1:2000) and goat anti-mouse antibody conjugated to horseradish peroxidase (KPL, 1:10,000) were used.

### 2.5 Statistical analysis

To determine the differences between sedentary and HIIT rats, unpaired t-tests were used to assess cocaine CPP, serum corticosterone and ΔFosB levels. Statistical significance for all tests was set to p<0.05.

## 3. Results

### 3.1 Cocaine Conditioned Place Preference

All data were analyzed via an unpaired t-test. Sedentary rats exhibited a preference for the cocaine-paired chamber, indicated by a significant increase in time spent in the cocaine chamber from Pretest to Test [t(14)= 2.833, p= 0.0133; Figure 2]. By contrast, the HIIT rats exhibited a place aversion to the cocaine-paired chamber, indicated by a significant decrease in time spent in the cocaine chamber from Pretest to Test [t(28)= 2.576, p= 0.0156; Figure 2]. Sedentary rats thus also showed a significantly increased amount of time spent in the cocaine chamber on Test Day compared to that of the HIIT subjects on Test Day [t(21)= 4.292, p= 0.0003; Figure 2].

**Fig 2.**
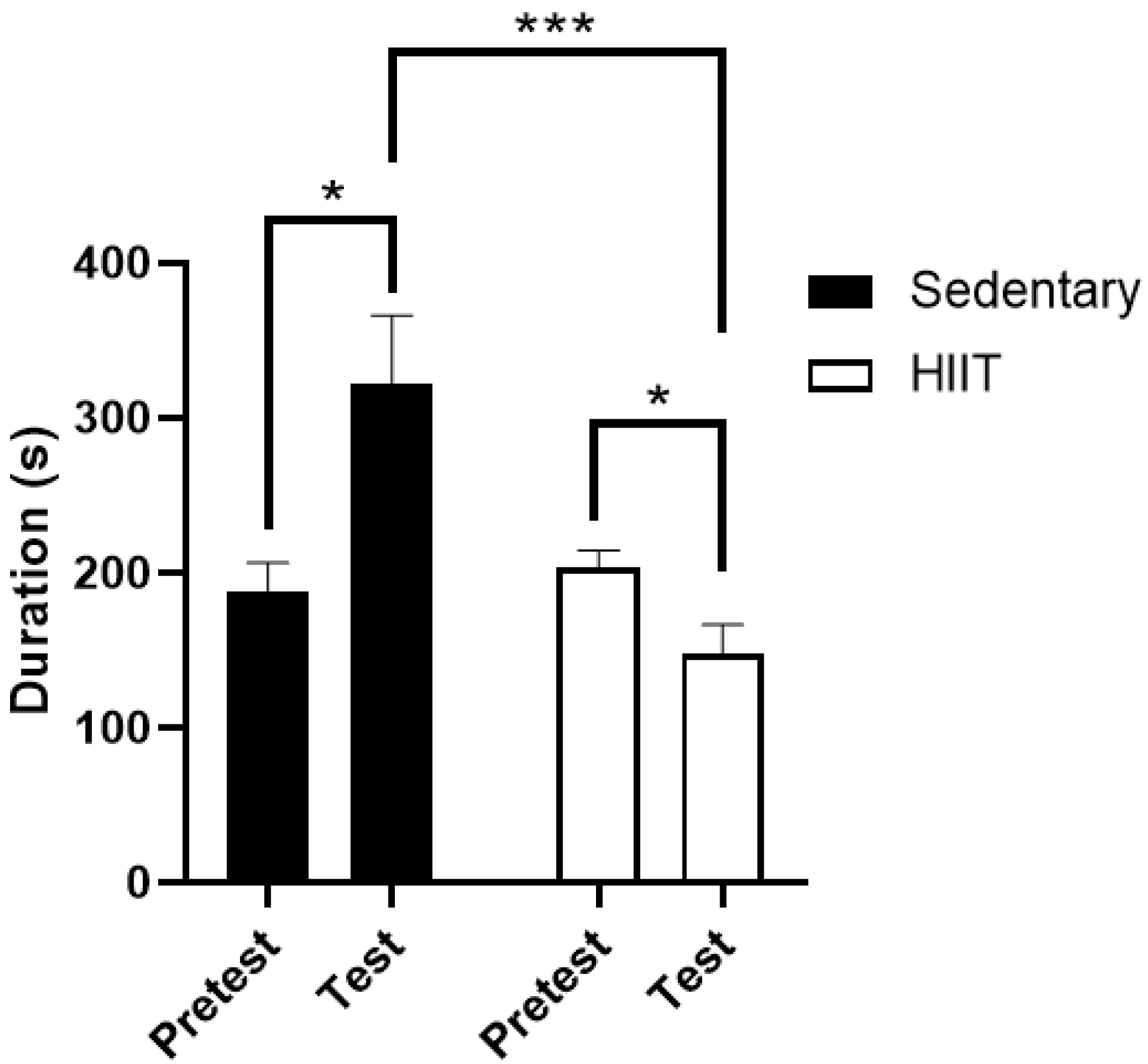
Mean time spent in cocaine chamber (sec + SEM) during Pretest and Test for both Sedentary and HIIT subjects. Sedentary subjects showed a significant preference for cocaine (*p ≤ 0.05). HIIT subjects showed a significant aversion to cocaine (*p ≤ 0.05). Sedentary subjects showed a significantly increased amount of time spent in the cocaine chamber on Test Day compared to that of HIIT subjects on Test Day (*p ≤ 0.001). Time spent in cocaine chamber during Pretest was not statistically significant between Sedentary and HIIT subjects.

### 3.2 Stress Reactivity Testing (Serum Corticosterone ELISA)

Both the HIIT and the sedentary group of rats were examined for serum corticosterone levels. A T-test revealed [t(10.42)= 0.5384, p= 0.6016] that there was no significant difference between the two groups in terms of serum corticosterone levels (p = ns; Figure 3).

**Fig 3.**
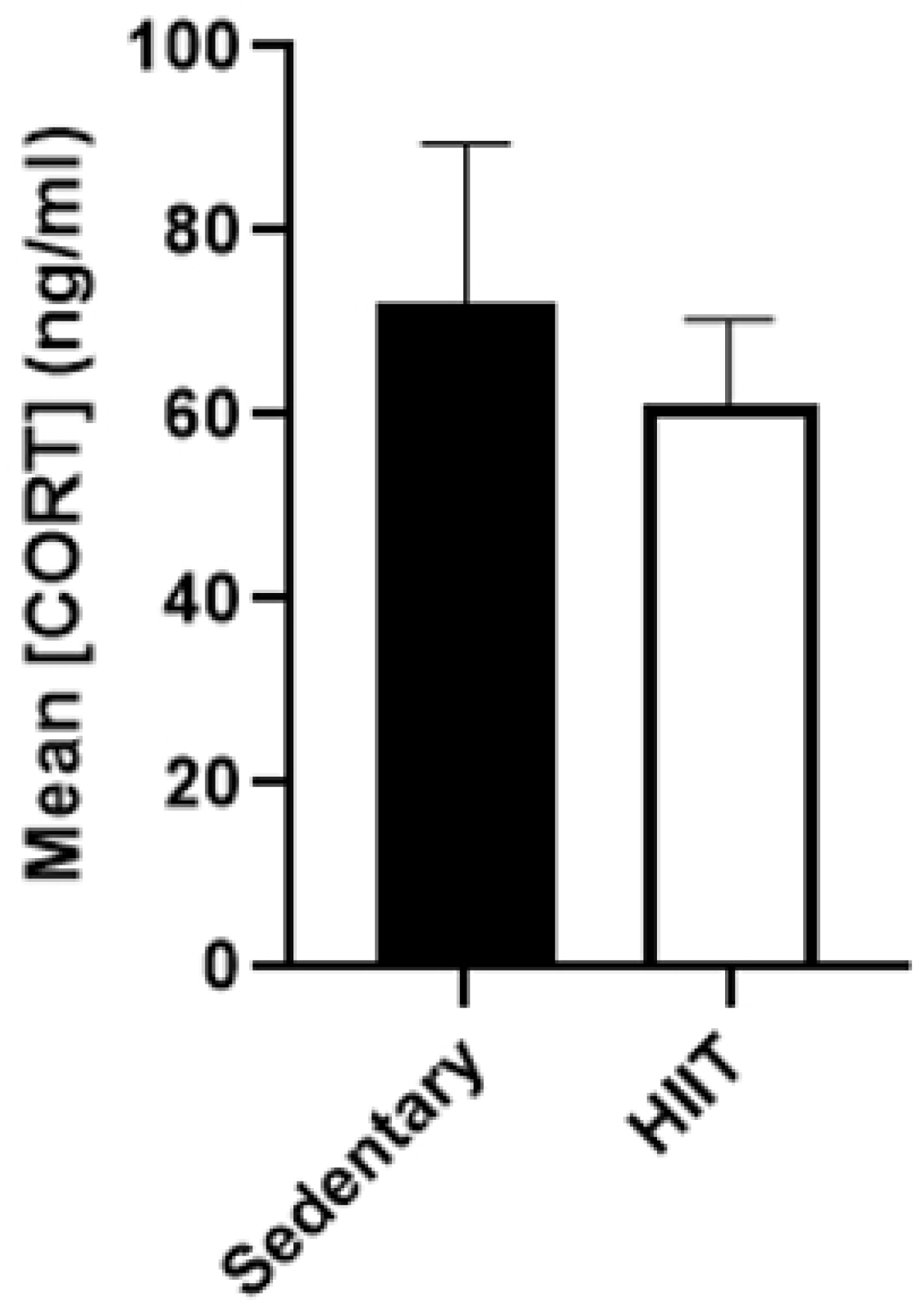
Mean corticosterone (ng/ML + SEM) for both sedentary and HIIT subjects following the chronic exercise regimen and prior to cocaine CPP. Both HIIT and Sedentary subjects showed similar levels of serum corticosterone. A t-test was ran and no significant difference was observed between the two groups (p= 0.6016).

### 3.3 Immunoblot Analysis (ΔFosB Western blot)

The dorsal and ventral striatum of the sedentary and HIIT treated rats were analyzed for ΔFosB levels. After running an unpaired t-test it was determined that there was no significant difference in ΔFosB levels between the dorsal and ventral striatum of the sedentary group [t(10)= 1.441, p= 0.1801]. There was also no significant different between the dorsal and ventral striatum of the HIIT group [t(15)= 1.672, p= 0.1153]. Due to this, we combined the dorsal and ventral striatum results and labeled them as striatum. There was a significant increase of ΔFosB in the HIIT compared to the sedentary group [t(28)= 2.184, p= 0.0375; Figure 4].

**Fig 4.**
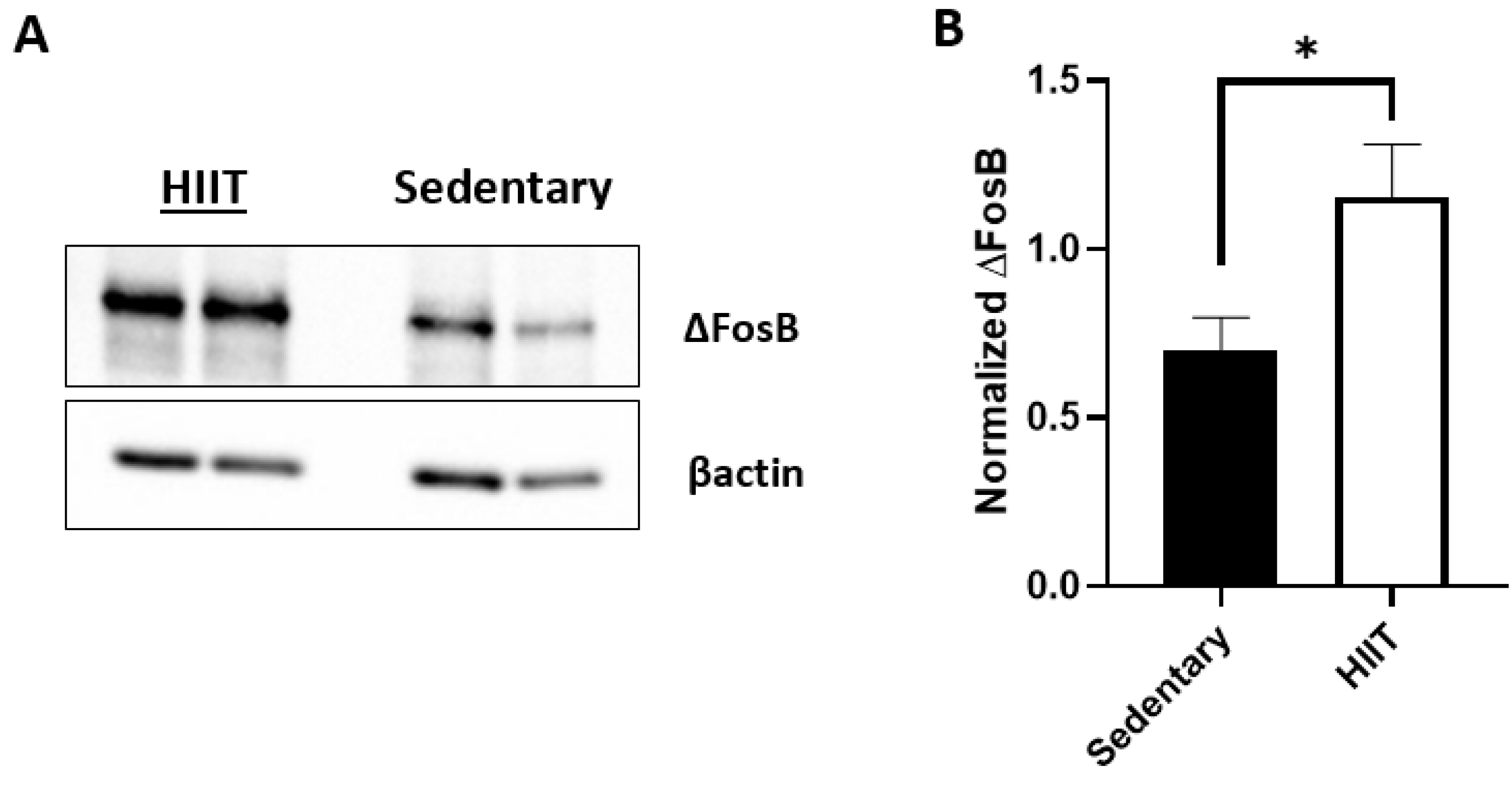
Western blot results of ΔFosB. A) Exemplary western blots of ΔFosB. There are two samples per regimen represented in this figure. FosB anti-rabbit antibody was used to detect ΔFosB in the striatal samples of the sedentary and HIIT animals. B) Normalized ΔFosB (+SEM) for both sedentary (n=12) and HIIT (n=18) following the chronic exercise regimen and cocaine CPP. The ΔFosB band was normalized to its corresponding βactin band. The HIIT subjects had significantly higher ΔFosB levels than the sedentary subjects (*p ≤ 0.05).

## 4. Discussion

The present study showed that rats exposed to a chronic HIIT exercise regimen displayed an aversion to cocaine as measured by the cocaine conditioned place preference paradigm. Our lab has previously reported the efficacy of treadmill exercise in decreasing cocaine preference [7]; however, this study exposed rats to chronic moderate-intensity continuous exercise (MICT), commonly regarded as standard aerobic exercise, and showed that exercise-treated rats still displayed a preference for cocaine, yet to a lesser degree. The present study builds on our previous findings by illustrating that HIIT may serve as a more effective intervention in the realm of cocaine abuse than that of MICT by not only decreasing cocaine preference but causing an aversion to cocaine. With these findings, the question arises as to what physiological and neurological mechanisms underlie the increased efficacy of HIIT compared to MICT relating to cocaine abuse.

In addressing this question, the first matter to be discussed is the differences between the effects of HIIT and MICT; here, a few general points of difference have been noted. When compared to MICT, HIIT has been shown to result in increased glucose metabolism [24], greater reductions in fat mass [25], and greater improvements in VO_2_ max [10], the lattermost of which is widely used as a measure of cardiorespiratory fitness [26]. HIIT has also been shown to possess enhanced efficacy in disease treatment: this phenomenon has been demonstrated with regards to cardiac disease [27–29], multiple sclerosis [30], and diabetes [31].

The other major matter to be discussed is that of general factors which affect cocaine CPP. It has been shown that the cannabinoid receptor system is implicated in cocaine abuse, with CB1 receptor antagonism and CB2 receptor agonism resulting in reduced cocaine preference [32]. It has previously been shown that MICT does not alter CB1 receptor levels in rats [33]; however, the effects of HIIT on CB1 and CB2 receptor levels has not been explored. It has been shown that the administration of THC increases Fos accumulation in cocaine exposed adolescent rats [34]. As a result, further investigation should be performed to expand upon the endocannabinoid system’s mediation in cocaine preference. Additionally, it has been shown that a neurotensin analog blocks cocaine CPP [35]. Neurotensin is a neuropeptide that has been strongly implicated to act with the dopaminergic system. Previous studies have shown that MICT decreased dopamine D1 receptor levels and increased D2 receptor levels, but this phenomenon has similarly not been explored in HIIT [33,36].

Aside from the aspect of exercise intensity, the role of sex has not been adequately examined with respect to HIIT. Our previous study [7] showed that MICT decreased cocaine preference in females while eliminating cocaine preference in males. In previous studies, 25 mg/kg of cocaine has induced a place preference [7,23]. The present study shows that HIIT not only eliminated cocaine preference in males but caused an aversion to the cocaine-paired environment. As a result, future studies should explore the efficacy of HIIT on cocaine preference in female rats, as our previous findings illustrate sex as a crucial factor. This occurrence is also supported by the literature, as females have been noted to possess higher levels of vulnerability than males during various phases of the addiction process, including acquisition, maintenance, and relapse [37]. According to Orihuel *et al.* [34] there were significant sex-dependent interactions between cocaine and adolescent THC exposure in the dorsal hypothalamus, suggesting that cocaine induced a more robust cellular activation in THC exposed females than males. Additionally, it has been shown that when given free access to a running wheel, female rats run significantly more than males [38], thus illustrating a sex difference with regards to exercise. As these factors point to sex differences regarding both exercise and addiction, they highlight the potential value in exploring sex as a factor in the role of exercise on addiction.

HIIT exercise pretreated rats tested for cocaine CPP show higher levels of ΔFosB in their striatum compared to their sedentary animal counterparts. Due to the sub-chronic nature of the cocaine injections, this increase in ΔFosB level was caused mostly by the chronic HIIT exercise regimen. Previous studies have shown that when a drug of abuse is administered, there is no increase in ΔFosB level without the functional D1 receptor [39–41]. This indicates that the D1 receptor is needed for ΔFosB expression. HIIT exercise when administered has demonstrated an increase of dopamine type 2-like receptor (D2R) binding in the nucleus accumbens, which has previously been linked to attenuating drug seeking behavior [42]. Males have shown higher levels of D2R binding after HIIT compared to females, making the females more susceptible to addiction [42]. Within the same study there was no significant difference seen in tyrosine hydroxylase or D1 receptor, an explanation for this was possible upper limits to neurotrophic factors due to the healthy nature of the animal subjects [42]. Due to the important role of dopamine signaling in addiction and exercise there is a need for further investigation on how HIIT affects the expression of the D1, D2 and D3 receptors.

As animals exercise, dopamine and ΔFosB levels increase [43,44]. Since ΔFosB has a long half-life [18,45], it is possible that the increase we saw due to HIIT exercise is attenuating or reversing the rewarding dopaminergic effects of cocaine place preference [46, 47]. In Zhang *et al.* [48], rats exposed to an enriched environment had higher baseline levels of ΔFosB than their counterparts. After cocaine self-administration, the enriched environment rats did not have a significant increase in ΔFosB. In addition, these rats had reduced cocaine-seeking behavior [48, 49]. Since exercise is considered a form of environmental enrichment [50,51], an increase in dopamine from HIIT could be blocking the rewarding sensation of cocaine, thus causing an aversion to the cocaine-paired chamber after undergoing chronic HIIT.

## 5. Conclusion

In conclusion, the present study showed that HIIT exercise during adolescence effectively prevented cocaine preference compared to control rats. Specifically, HIIT exercise in adolescent rats produced a significant aversion to the cocaine-paired environment. These results encourage and further support the benefit of aerobic exercise, and in particular HIIT, in reducing the risk of substance-related behaviors, such as cocaine preference. Future research will further explore the underlying mechanisms behind this phenomenon of HIIT exercise on ΔFosB and substance abuse. Finally, these findings support the concept of specific exercise dosing regimens like HIIT having distinct effects on drug abuse behavior mediated by ΔFosB and could have important future implications for a personalized medicine approach to drug abuse intervention.

## Acknowledgements

This research was funded by the New York Research Foundation #RIAQ0094. We thank Sierra Douglas for help with animal handling and behavioral testing.

